# Serum Calprotectin as predictor of Rheumatoid Arthritis disease severity and Mortality

**DOI:** 10.64898/2026.10.30.763141

**Authors:** Seimia Chebbi, Denis Mongin, Delphine Courvoisier, Michael J Nissen, Burkhard Möller, Raphael Micheroli, Cem Gabay, Nicolas Vuilleumier, Axel Finckh, Matthias Jarlborg

**Affiliations:** Division of Laboratory Medicine, Diagnostic Department, Geneva University Hospitals, Geneva, Switzerland; Department of Medicine, Medical Faculty, Geneva University, Geneva, Switzerland. Division of Rheumatology, University Hospitals of Geneva, CH-1206 Geneva, Switzerland; Rheumatology, Immunology and Allergy, Inselspital, University Hospital of Bern, CH-3010 Bern, Switzerland; SCQM Biobank Study Group, SCQM Registry, Foundation SCQM, Zürich, Switzerland; Department of Rheumatology, University Hospital of Zurich, University of Zurich, Switzerland; Geneva Center of Inflammation Research, University of Geneva Faculty of Medicine, Geneva, Switzerland

## Abstract

**Objective:** To evaluate the clinical utility of serum calprotectin (sCAL) as a biomarker of disease activity, disease severity, cardiovascular risk, and mortality in inflammatory rheumatic diseases (IRDs).

**Methods:** Serum calprotectin was measured in 2677 patients with IRDs, including 1472 rheumatoid arthritis (RA), 771 axial spondyloarthritis (axSpA), and 434 psoriatic arthritis (PsA) patients enrolled in the Swiss Clinical Quality Management in Rheumatic Diseases registry, as well as 248 healthy blood donors. Associations with disease activity, radiographic progression, cardiovascular biomarkers, incident major adverse cardiovascular events (MACE), and mortality were evaluated using receiver operating characteristic, Cox regression, and Kaplan-Meier models.

**Results:** sCAL concentrations were significantly higher in RA than in axSpA, PsA, or healthy blood donors. Although diagnostic discrimination between IRDs was modest (Area under the curve (AUC): 0.56), elevated sCAL identified a subset of patients with RA characterized by higher disease activity, higher autoantibody levels, and accelerated radiographic progression, performing better than CRP for the assessment of inflammatory activity, particularly for the discrimination of active synovitis (SJC28≥4, AUC 0.67 vs 0.62, p=0.012), even in patients with normal CRP concentrations. Elevated sCAL was independently associated with increased all-cause mortality (HR:2.99; 95% confidence interval (95%CI):1.87-4.78), cardiovascular mortality (HR:7.49;95%CI:1.50-37.55), respiratory mortality (HR: 5.89; 95%CI:1.97-17.62), and incident MACE (HR: 2.10; 95%CI: 1.04-4.24). Furthermore, sCAL correlated with high-sensitivity cardiac troponin T and NT-proBNP concentrations, independently of lipid profile.

**Conclusions:** Serum calprotectin is a clinically relevant biomarker in RA that reflects both disease activity and severity, independently identifying patients at increased cardiovascular risk and mortality, supporting its potential role in disease monitoring and risk stratification.

## Introduction

Inflammatory rheumatic diseases (IRDs) are a heterogeneous group of immune-mediated disorders characterized by persistent joint inflammation, progressive structural damage, and significant functional impairment. Together, they impose a considerable global burden, affecting more than 5% of the population [1]. Rheumatoid arthritis (RA), axial spondyloarthritis (axSpA), and psoriatic arthritis (PsA) account for the majority of cases and share key pathogenic features, notably systemic inflammation and synovitis [2]. Accurate assessment of inflammation is central for diagnosis, disease monitoring and therapeutic response assessment. In clinical practice, the acute-phase response - most commonly measured by erythrocyte sedimentation rate (ESR) and C-reactive protein (CRP) - remains the cornerstone of inflammation quantification [3] [4]. However, these biomarkers have important limitations. Their levels display marked heterogeneity across patient populations and disease entities; and targeted therapies, particularly interleukin-6 (IL-6) inhibitors, may suppress their expression independently of clinical response. As a result, reliance on conventional acute-phase reactants carries the risk of underestimating disease activity in a substantial proportion of patients [5] [6] , or conversely, of overestimating it, as ESR may be elevated due to comorbidities such as anemia or dysproteinemia associated with Sjögren syndrome.

Identifying biomarkers that reflect more directly the underlying inflammatory processes within affected tissues is crucial [7]. Synovitis, the pathological hallmark of IRDs, is characterized by the infiltration and activation of immune cells, with granulocytes representing a major cellular component of the synovial fluid and inflamed synovium [8] [9]. Granulocyte-derived alarmins have therefore emerged as promising candidates for capturing local inflammatory activity [10]. Calprotectin, a heterodimer of S100A8 and S100A9, is abundantly expressed in the cytosol of granulocytes and released upon cellular activation or damage [12]. Owing to its small molecular size and remarkable stability, it readily diffuses into the circulation, where it can be reliably quantified [13]. Importantly, serum calprotectin has been shown to correlate with inflammatory activity at the tissue level, suggesting that it may provide a more sensitive and direct measure of ongoing synovitis than conventional acute-phase reactants [11].

Beyond its role as a marker of disease activity, emerging evidence suggests that serum calprotectin (sCAL) may also reflect systemic inflammatory burden more broadly and could be associated with clinically relevant outcomes, including mortality and cardiovascular (CV) risk in patients with IRDs. However, the prognostic value of sCAL remains insufficiently characterized.

The present study therefore aims to evaluate the utility of sCAL as a biomarker of disease activity and severity in inflammatory rheumatic diseases. In addition, we seek to assess its potential role as a prognostic biomarker for mortality and cardiovascular risk, and to determine whether it could serve as a reliable alternative to conventional acute-phase reactants such as CRP in contemporary therapeutic settings.

## Samples and Methods

### Study populations The SCQM registry

The Swiss Clinical Quality Management in Rheumatic Diseases (SCQM) registry is a nationwide, prospective registry for patients with IRDs (www.scqm.ch), currently including over 10959 patients with RA, 5875 with axSpA, and 3425 with PsA, with data collection starting in 1997, 2005, 2006, respectively. The median year of inclusion was 2010 (IQR 2004–2015) for RA, 2014 (IQR 2009–2019) for axSpA, 2015 (IQR 2010–2020) for PsA, and 2022 (IQR 2021–2023) [14]. Serum samples were obtained from patients enrolled until 2021 with established diagnoses of RA (1472), axSpA (771), and PsA (434). Briefly, founded in 1997 the SCQM registry enrolls patients originating from private practices (50%), academic centers (20%), and non-academic centers (30%), providing a real-life IRD population sample of Switzerland. Follow-up data collection, including cardiovascular events and associated risk factors encompassing the period before and after inclusion in the registry, is provided by treating rheumatologists through dedicated case report forms. Regular assessments are performed, approximately 1–2 times per year (including disease activity, anti-rheumatic treatments, side effects, reasons for discontinuation, and comorbidities).

Clinical, laboratory, treatment, and outcome data were extracted from the SCQM registry according to predefined study endpoints (see below) by data scientists blinded to biochemical results. The study was approved by the local ethics committee of the University Hospital of Geneva, the SCQM Biobank Scientific Advisory Board, and SCQM Foundation board (PB_2018-00317) and abides by the principles of Helsinki declaration.

### Healthy blood donors

Two-hundred and forty-eight healthy blood donors (122 women and 126 men; age range 18–74 years) were recruited under the standards ISO 1589/2022 and Clinical and Laboratory Standard Institute (CLSI) EP28-A3c guidelines to derive and/or validate 95% reference intervals on different biological matrixes [15]. Accordingly, the upper limit of the reference interval for serum calprotectin was defined as the 97.5th percentile of the healthy donor distribution. Based on this analysis, a pre-specified cutoff of 4.2 μg/mL was selected to define elevated sCAL concentrations and further validated in post-hoc receiver operating characteristic (ROC) analyses (Supplementary figure 1). Patients were subsequently classified into high- and low-sCAL groups according to this threshold.

### Study design

Initially, we evaluated the biosamples in the SCQM registry to determine sCAL differences between various IRDs subgroups and healthy blood donors. In a second step we examined sCAL as a biomarker of disease activity, disease severity, and long-term outcomes in IRDs. The biomarker of interest was serum calprotectin, with CRP serving as the comparator biomarker.

### Biomarker measurements

Serum calprotectin concentrations were measured using the BÜHLMANN sCAL® turbo assay (BÜHLMANN Laboratories AG, Schönenbuch, Switzerland), an IVDR-certified particle-enhanced immunoturbidimetric assay performed on a cobas c502 analyzer (Roche Diagnostics, Basel, Switzerland) in the Laboratory Medicine Service of Geneva University Hospitals. Assays were performed according to the manufacturer’s instructions and ISO 1589/2022 procedures. CRP, high-density lipoprotein (HDL) cholesterol, low-density lipoprotein (LDL) cholesterol, and triglycerides were measured on the same analyzer, while high-sensitivity cardiac troponin T (hs-cTnT), N-terminal pro-B-type natriuretic peptide (NT-proBNP) levels (cardiac biomarkers) were measured on cobas 802 analysers (Roche Diagnostics, Basel, Switzerland) according to the manufacturer’s instructions and ISO 1589/2022 procedures. A CRP concentration ≥5 mg/L was considered elevated according to institutional reference standards.

### Clinical outcomes

The primary outcome of interest was all-cause mortality, with disease activity and severity as factors associated with mortality. The occurrence of MACE and mortality was assessed using longitudinal follow-up data. Follow-up was based on the date of serum sampling, which served as baseline (time zero). Participants were followed until the occurrence of the outcome of interest or censoring. All-cause mortality, cardiovascular mortality, and cause-specific mortality were analyzed. Cause-specific categories included respiratory disease, infection, cancer, dementia, accidental death, and other causes.

Disease activity was assessed using swollen joint counts (SJC) and tender joint counts (TJC) across all disease groups. Disease-specific activity measures included the Clinical Disease Activity Index (CDAI), Disease Activity Score in 28 joints (DAS28), patient global assessment (PGA), and physician global assessment (PhGA) for RA; the Bath Ankylosing Spondylitis Functional Index (BASFI), Bath Ankylosing Spondylitis Disease Activity Index (BASDAI), and Ankylosing Spondylitis Disease Activity Score (ASDAS) for axSpA; and the Disease Activity Index for Psoriatic Arthritis (DAPSA) and measures of skin involvement for PsA. Markers of disease severity and prognostic outcomes included autoantibody status, cardiovascular risk biomarkers, radiographic damage progression assessed incident and prevalent major adverse cardiovascular events (MACE), and all-cause and cause-specific mortality. In patients with RA, longitudinal radiographic progression assessed by increasing Ratingen damage score on sequential radiographs, and changes in disease activity were additionally evaluated.

### Statistical analysis

Continuous variables are presented as median (interquartile range) or mean (standard deviation), as appropriate. Categorical variables are expressed as counts and percentages. Because serum calprotectin concentrations exhibited a non-normal distribution (right-skewed) and heteroscedasticity across groups, between-group comparisons were performed using Kruskal–Wallis tests followed by Dunn’s post hoc correction for multiple comparisons, with false discovery rate (FDR) control using the Benjamini–Hochberg method.

Pairwise comparisons were performed using RA as the reference group. Correlations between biomarkers and clinical outcomes were assessed using Spearman’s rank correlation coefficients. Receiver operating characteristic (ROC) analyses were performed to evaluate the discriminative performance of sCAL and CRP. Areas under the ROC curves (AUCs) were compared using DeLong’s test. Time-to-event analyses were performed using Kaplan–Meier survival curves and Cox proportional hazards regression models. Hazard ratios (HRs) and 95% confidence intervals (CIs) were estimated before and after adjustment for predefined covariates, including age, sex, disease duration, and additional confounders where indicated. Logistic regression analyses were used to assess independent associations between elevated sCAL and cardiovascular biomarkers. All statistical tests were two-sided, and p-values <0.05 were considered statistically significant. Analyses were performed using R software (R Foundation for Statistical Computing, Vienna, Austria).

## Results

### Serum calprotectin is elevated in RA compared with HBD and other inflammatory rheumatic diseases

Baseline characteristics of the study cohort are summarized in Table 1. Among the 2677 patients included, 1472 had RA, 771 AS, and 434 PsA. Patients with RA were older and more frequently female than those with axSpA or PsA (all p<0.001). RA patients also exhibited a higher prevalence of hypertension, diabetes, and prevalent MACE. Consistent with this cardiovascular burden, circulating hs-cTnT and NT-proBNP concentrations were significantly higher in RA compared to axSpA or PsA (all p<0.001). sCAL concentrations differed significantly across disease groups and were highest in RA (1.70 [1.00–2.80] μg/mL), compared with axSpA (1.50 [1.00, 2.40]) and PsA (1.40 [0.90, 2.10], p<0.001; Figure 1A). In contrast, CRP levels were comparable across diseases and did not differ significantly between groups (p=0.066; Table 1), highlighting the differential sensitivity of these biomarkers. Although sCAL concentrations were elevated in RA, its ability to discriminate RA from AS or PsA remained modest, with an area under the receiver operating characteristic curve (AUC) of 0.56. Nevertheless, sCAL outperformed CRP for disease discrimination (p=0.007, Delong test; Figure 1B). At the threshold of 4.2 μg/mL, sCAL demonstrated high specificity (94.3%) but low sensitivity (14.3%) for RA discrimination, with a positive predictive value of 75,5%. Overall, 209 patients with RA (14.2%) had sCAL concentrations above this threshold (median 6.30 [5.10-8.40] ug/mL), compared with 46 AS patients (6%, median 5.05, [4.7-5.77] ug/mL) and 22 PsA patients (5.1%, median 4.80 [4.45-6.20] ug/mL. Across disease groups, patients with elevated sCAL displayed distinct clinical characteristics. In RA, elevated sCAL was associated with male sex and current smoking status (Table 2). A similar association with smoking was observed in PsA but not in axSpA (Supplementary Tables 1–2). Taken together, these findings demonstrate that circulating calprotectin levels are preferentially elevated in RA compared with other inflammatory rheumatic diseases, despite comparable CRP concentrations.

**Figure 1.**
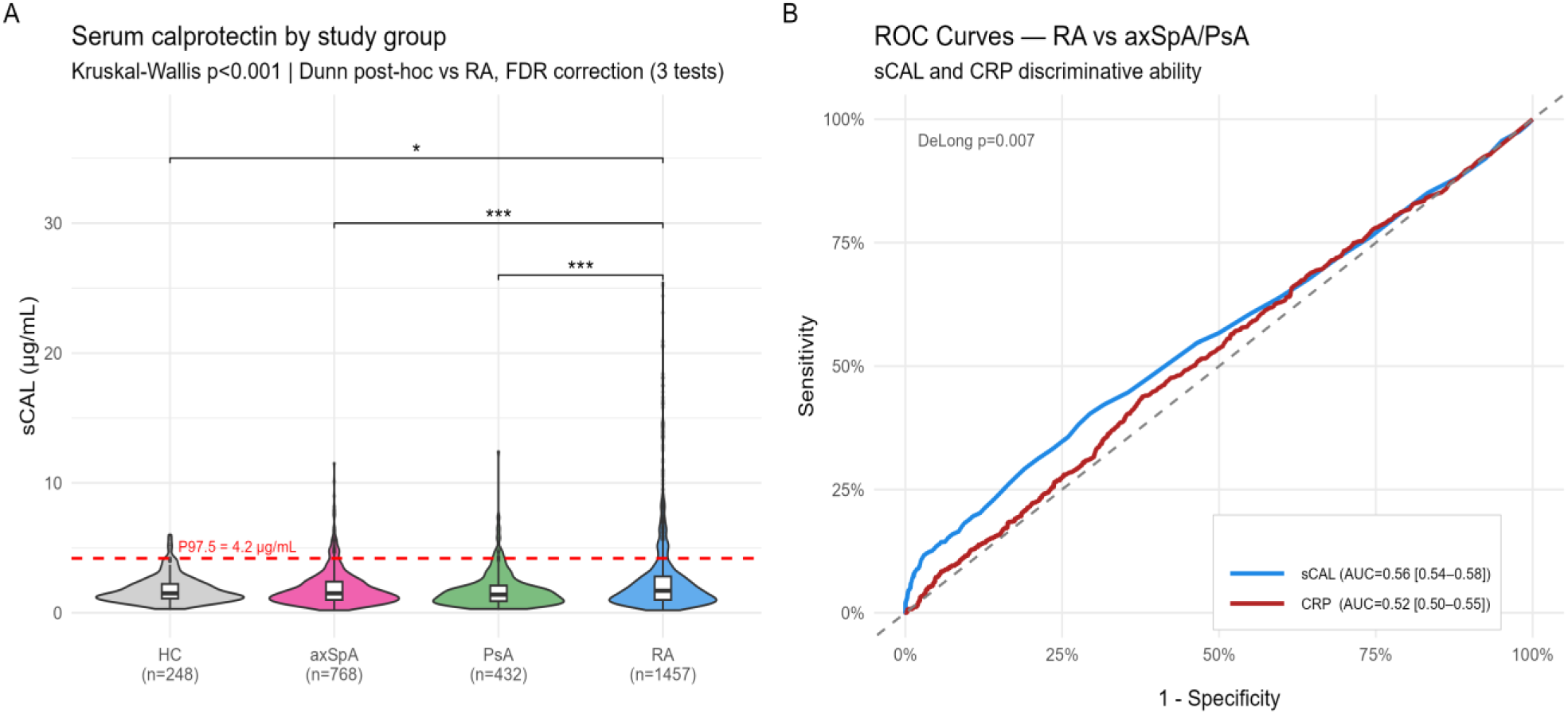
Serum calprotectin (sCAL) is elevated in rheumatoid arthritis (RA) compared to healthy controls (HC) and other inflammatory rheumatic diseases. (**A**) Distribution of sCAL levels in healthy controls and patients with RA, axial spondyloarthritis (axSpA), and psoriatic arthritis (PsA). The red dashed line indicates the upper reference limit (P97.5 = 4.2 µg/mL) established from a healthy donor reference cohort (n=248) in accordance with CLSI EP28-A3c guidelines. Boxes indicate median and interquartile range. Statistical comparisons were performed using the Kruskal-Wallis test followed by Dunn’s post-hoc test with false discovery rate correction (Benjamini-Hochberg, RA as reference, 3 comparisons). *p<0.05, ***p<0.001. (**B**) Receiver Operating Characteristic (ROC) curves for sCAL and CRP to discriminate RA from axSpA/PsA. AUC comparison by DeLong method.

**Table 1.** Baseline characteristics of the study cohort stratified by disease.

| Variable | Overall<br>(n=2,677) | RA<br>(n=1,472) | axSpA<br>(n=771) | PsA<br>(n=434) | p-value |
|---|---|---|---|---|---|
| **Demographics** |  |  |  |  |  |
| Age, years | 53.30 [42.60, 63.70] | 59.00 [49.88, 67.82] | 43.10 [34.70, 52.00] | 51.85 [43.15, 59.50] | <0.001 |
| Male sex, n (%) | 1069 (39.9) | 372 (25.3) | 452 (58.6) | 245 (56.5) | <0.001 |
| Smoker, n (%) | 975 (36.4) | 393 (26.7) | 462 (59.9) | 120 (27.6) | <0.001 |
| Disease duration, years | 6.88 [2.94, 13.58] | 7.75 [3.36, 14.68] | 6.46 [2.51, 12.84] | 5.29 [2.34, 10.80] | <0.001 |
| **Inflammatory biomarkers** |  |  |  |  |  |
| CRP, mg/L | 1.90 [0.75, 5.21] | 2.08 [0.79, 5.41] | 1.84 [0.67, 5.50] | 1.72 [0.75, 4.38] | 0.066 |
| sCAL, µg/mL | 1.60 [1.00, 2.50] | 1.70 [1.00, 2.80] | 1.50 [1.00, 2.40] | 1.40 [0.90, 2.10] | <0.001 |
| **Comorbidities** |  |  |  |  |  |
| Hypertension, n (%) | 691 (25.8) | 464 (31.5) | 125 (16.2) | 102 (23.5) | <0.001 |
| Diabetes, n (%) | 215 (8.0) | 136 (9.2) | 46 (6.0) | 33 (7.6) | 0.024 |
| **Cardiovascular biomarkers** |  |  |  |  |  |
| hs-cTnT, ng/L | 3.90 [0.00, 7.36] | 4.40 [0.00, 8.66] | 3.19 [0.00, 5.45] | 3.92 [0.00, 6.52] | <0.001 |
| NT-proBNP, pg/mL | 62.60 [33.70, 120.00] | 81.80 [45.20, 154.00] | 43.65 [25.10, 80.35] | 52.10 [30.23, 87.42] | <0.001 |
| HDL-cholesterol, mmol/L | 1.40 [1.13, 1.72] | 1.51 [1.23, 1.85] | 1.28 [1.07, 1.58] | 1.27 [1.06, 1.57] | <0.001 |
| LDL-cholesterol, mmol/L | 1.38 [0.97, 1.97] | 1.37 [0.97, 1.92] | 1.37 [0.94, 1.94] | 1.49 [1.08, 2.15] | 0.001 |
| Triglycerides, mmol/L | 2.98 [2.37, 3.64] | 3.00 [2.36, 3.66] | 2.84 [2.32, 3.49] | 3.09 [2.50, 3.79] | 0.001 |
| **Treatments** |  |  |  |  |  |
| Glucocorticoids, n (%) | 434 (16.2) | 400 (27.2) | 12 (1.6) | 22 (5.1) | <0.001 |
| csDMARDs, n (%) | 1599 (59.7) | 1101 (74.8) | 226 (29.3) | 272 (62.7) | <0.001 |
| Statins, n (%) | 357 (13.3) | 251 (17.1) | 62 (8.0) | 44 (10.1) | <0.001 |
| Prior bDMARDs, n (%) | 2316 (86.5) | 1270 (86.3) | 672 (87.2) | 374 (86.2) | 0.823 |
| Current bDMARD, n (%) |  |  |  |  | <0.001 |
| None | 857 (32.0) | 455 (30.9) | 254 (32.9) | 148 (34.1) |  |
| Anti-TNF | 1113 (41.6) | 390 (26.5) | 496 (64.3) | 227 (52.3) |  |
| Abatacept | 146 (5.5) | 142 (9.6) | 1 (0.1) | 3 (0.7) |  |
| Anti-IL6 | 174 (6.5) | 165 (11.2) | 5 (0.6) | 4 (0.9) |  |
| JAK inhibitor | 39 (1.5) | 36 (2.4) | 0 (0.0) | 3 (0.7) |  |
| Other | 348 (13.0) | 284 (19.3) | 15 (1.9) | 49 (11.3) |  |
| **Outcomes** |  |  |  |  |  |
| Prevalent MACE, n (%) | 179 (6.7) | 133 (9.0) | 27 (3.5) | 19 (4.4) | <0.001 |
| Incident MACE, n (%) | 47 (1.8) | 35 (2.4) | 5 (0.6) | 7 (1.6) | 0.012 |
| All-cause death, n (%) | 90 (3.4) | 78 (5.3) | 5 (0.6) | 7 (1.6) | <0.001 |

*Data are presented as median [interquartile range] for continuous variables and n (%) for categorical variables. p-values: Kruskal-Wallis test for continuous variables; Chi-squared or Fisher's exact test for categorical variables.*
*Abbreviation: axSpA, axial spondyloarthritis; bDMARD, biologic disease-modifying antirheumatic drug; CRP, C-reactive protein; csDMARD, conventional synthetic disease-modifying antirheumatic drug; HDL, high-density lipoprotein; hs-cTnT, high-sensitivity cardiac troponin T; LDL, low-density lipoprotein; MACE, major adverse cardiovascular event; NT-proBNP, N-terminal pro-B-type natriuretic peptide; PsA, psoriatic arthritis; RA, rheumatoid arthritis; sCAL, serum calprotectin.*

**Table 2.** Baseline characteristics of RA patients stratified by sCAL status.

| Variable | Overall<br>(n=1,472) | Low sCAL<br>(<4.2 µg/mL, n=1,263) | High sCAL<br>(>=4.2 µg/mL, n=209) | p-value |
|---|---|---|---|---|
| **Demographics** |  |  |  |  |
| Age, years | 59.00 [49.88, 67.82] | 58.85 [49.70, 67.70] | 59.10 [50.40, 68.10] | 0.590 |
| Male sex, n (%) | 372 (25.3) | 291 (23.3) | 76 (36.4) | <0.001 |
| Smoker, n (%) | 393 (26.7) | 311 (24.9) | 82 (39.2) | <0.001 |
| Disease duration, years | 7.75 [3.36, 14.68] | 7.74 [3.38, 14.56] | 7.93 [3.34, 15.11] | 0.826 |
| **Inflammatory biomarkers** |  |  |  |  |
| CRP, mg/L | 2.08 [0.79, 5.41] | 1.71 [0.72, 4.26] | 7.90 [2.90, 21.72] | <0.001 |
| sCAL, µg/mL | 1.70 [1.00, 2.80] | 1.40 [0.90, 2.20] | 6.30 [5.10, 8.40] | <0.001 |
| **Disease activity** |  |  |  |  |
| DAS28 | 2.60 [1.80, 3.80] | 2.50 [1.70, 3.60] | 3.90 [2.60, 4.90] | <0.001 |
| CDAI | 8.00 [3.00, 16.00] | 7.00 [3.00, 14.00] | 11.50 [5.00, 24.00] | <0.001 |
| Swollen joint count (28) | 0.00 [0.00, 3.00] | 0.00 [0.00, 2.00] | 2.00 [0.00, 7.00] | <0.001 |
| Tender joint count (28) | 1.00 [0.00, 4.00] | 1.00 [0.00, 4.00] | 2.00 [0.00, 6.00] | <0.001 |
| **Serology\*** |  |  |  |  |
| Seropositive, n (%) | 1050 (78.1) | 864 (75.9) | 175 (90.7) | <0.001 |
| RF positive, n (%) | 944 (70.5) | 770 (67.9) | 165 (85.5) | <0.001 |
| Anti-CCP positive, n (%) | 895 (68.5) | 730 (66.0) | 155 (82.4) | <0.001 |
| **Comorbidities** |  |  |  |  |
| Hypertension, n (%) | 464 (31.5) | 388 (31.1) | 71 (34.0) | 0.454 |
| Diabetes, n (%) | 136 (9.2) | 114 (9.1) | 21 (10.0) | 0.770 |
| **Cardiovascular biomarkers** |  |  |  |  |
| hs-cTnT, ng/L | 4.40 [0.00, 8.66] | 4.23 [0.00, 8.27] | 5.30 [3.06, 10.40] | 0.001 |
| NT-proBNP, pg/mL | 81.80 [45.20, 154.00] | 78.70 [44.38, 146.00] | 97.60 [50.50, 182.00] | 0.008 |
| HDL-cholesterol, g/L | 1.51 [1.23, 1.85] | 1.52 [1.23, 1.85] | 1.43 [1.20, 1.75] | 0.034 |
| **Treatments** |  |  |  |  |
| Glucocorticoids, n (%) | 400 (27.2) | 313 (25.1) | 82 (39.2) | <0.001 |
| csDMARDs, n (%) | 1101 (74.8) | 932 (74.7) | 156 (74.6) | 1.000 |
| Statins, n (%) | 251 (17.1) | 205 (16.4) | 44 (21.1) | 0.122 |
| Prior bDMARDs, n (%) | 1270 (86.3) | 1072 (85.9) | 188 (90.0) | 0.140 |
| Current bDMARD, n (%) |  |  |  | 0.023 |
| None | 455 (30.9) | 381 (30.5) | 65 (31.1) |  |
| Anti-TNF | 390 (26.5) | 351 (28.1) | 37 (17.7) |  |
| Abatacept | 142 (9.6) | 118 (9.5) | 24 (11.5) |  |
| Anti-IL6 | 165 (11.2) | 139 (11.1) | 25 (12.0) |  |
| JAK inhibitor | 36 (2.4) | 26 (2.1) | 8 (3.8) |  |
| Other | 284 (19.3) | 233 (18.7) | 50 (23.9) |  |
| **Outcomes** |  |  |  |  |
| Prevalent MACE, n (%) | 133 (9.0) | 108 (8.7) | 24 (11.5) | 0.235 |
| Incident MACE, n (%) | 35 (2.4) | 26 (2.1) | 9 (4.3) | 0.089 |
| All-cause death, n (%) | 78 (5.3) | 53 (4.2) | 23 (11.0) | <0.001 |

*Data are presented as median [interquartile range] for continuous variables and n (%) for categorical variables. p-values: Wilcoxon rank-sum test (continuous); Chi-squared or Fisher's exact test (categorical).*
*\*Serology data available in a subset of RA patients.*
*Abbreviation: Anti-CCP, anti-cyclic citrullinated peptide; bDMARD, biologic disease-modifying antirheumatic drug; CDAI, Clinical Disease Activity Index; CRP, C-reactive protein; csDMARD, conventional synthetic DMARD; DAS28, Disease Activity Score 28 joints; HDL, high-density lipoprotein; hs-cTnT, high-sensitivity cardiac troponin T; LDL, low-density lipoprotein; MACE, major adverse cardiovascular event; NT-proBNP, N-terminal pro-B-type natriuretic peptide; PSA, psoriatic arthritis; RA, rheumatoid arthritis; sCAL, serum calprotectin.*

### Serum calprotectin correlates with joint inflammation in rheumatoid arthritis

Given the preferential elevation of sCAL in RA, we investigated its relationship with disease activity and joint inflammation. In patients with RA, sCAL levels were positively correlated with SJC (r = 0.27, p < 0.001), indicating an association between circulating calprotectin and clinically apparent synovitis (Figure 2A). Consistent with this finding, sCAL also correlated significantly with TJC (r=0.19, p<0.001), as well as with composite disease activity indices, including CDAI (r=0,22, p<0.001) and the DAS28 (r=0.32, p<0.001; Figure 2B). Notably, sCAL outperformed the CRP for the assessment of joint counts and CDAI (SJC28: r=0.27 versus 0.22; TJC28: r=0.19 versus 0.16; CDAI: r=0,22 versus 0.18) whereas CRP showed a stronger correlation with DAS28 (r=0.41 versus 0.32), likely reflecting the mathematical incorporation of CRP into this composite score. Overall, these findings suggest that sCAL may more closely reflect articular inflammation in RA than CRP. The association between sCAL and SJC was also observed in PsA (r=0.18, p=0.016, Figure 2D), another disease characterized by peripheral joint involvement, although the correlations were weaker than those observed in RA. In contrast, no significant association was observed between sCAL and SJC in ankylosing spondylitis (axSpA, figure 2C), a disease predominantly characterized by axial inflammation. Nevertheless, sCAL demonstrated significant correlations with composite measures of axial disease activity and functional impairment, including the BASDAI (r=0.11, p=0.008), the ASDAS (r=0.17, p<0.001), and the BASFI (r=0.09, p=0.016). However, these correlations were generally weaker than associations with disease activity measures for RA. Taken together, these findings suggest that sCAL is most strongly associated with clinical disease activity in RA, particularly with measures reflecting peripheral joint inflammation. While its associations in axSpA and PsA, though statistically significant, were generally weaker and comparable to or inferior to those observed with CRP.

**Figure 2.**
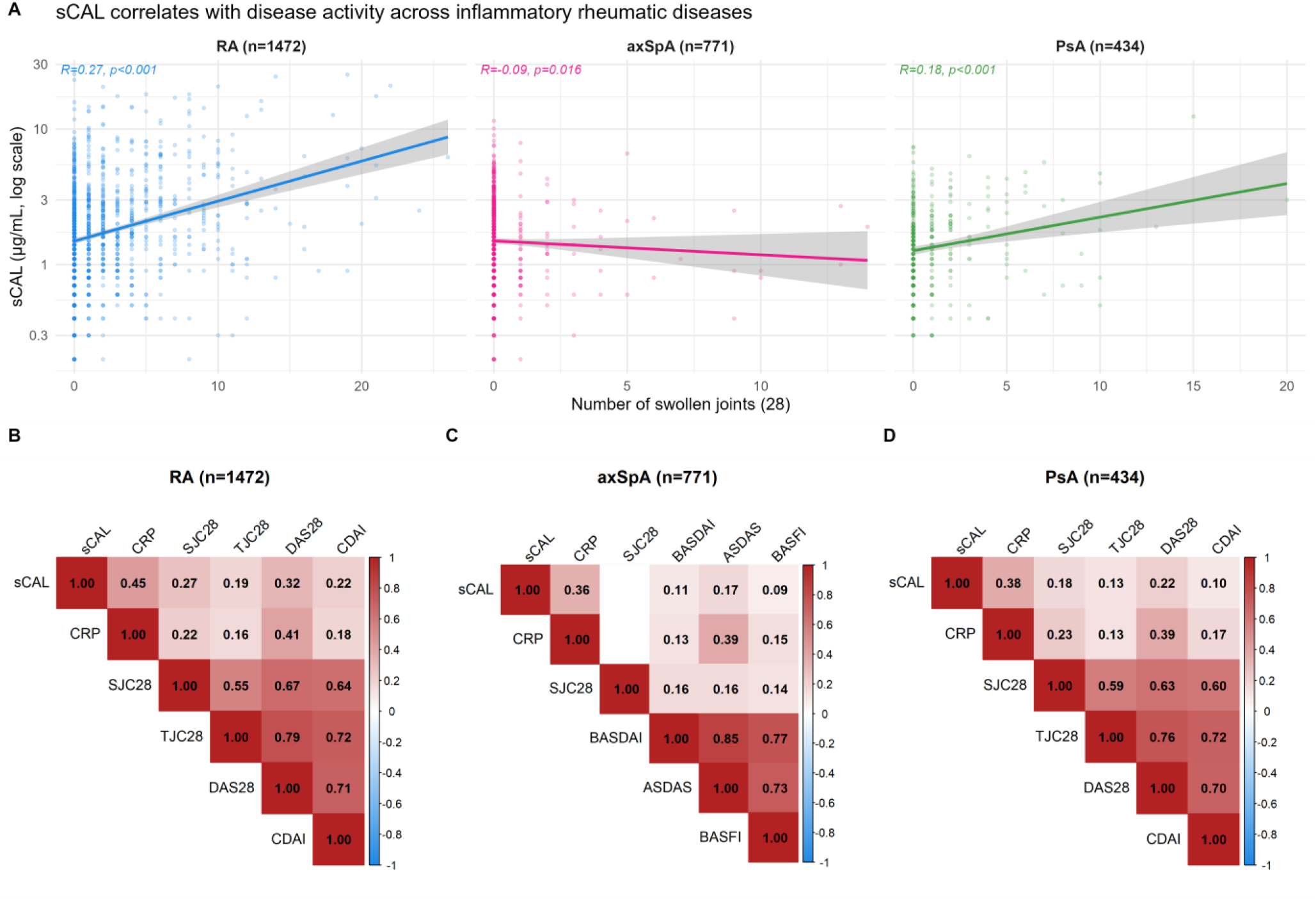
sCAL correlates with markers of disease activity across inflammatory rheumatic diseases. **(A)** Association between swollen joint count (SJC28) and sCAL levels by disease group. Lines represent linear regression fits with 95% confidence intervals. Spearman correlation coefficient (R) and p-value are shown. **(B-D)** Spearman correlation matrices between sCAL, CRP and disease activity scores in RA (B), axSpA (C), and PsA (D). Color gradient indicates the strength and direction of Spearman correlations, ranging from red (positive correlation) to blue (negative correlation). Numbers indicate the Spearman correlation coefficient. Blank cells indicate non-significant correlations (p≥0.05). **Abbreviations**: ASDAS, Ankylosing Spondylitis Disease Activity Score; BASDAI, Bath Ankylosing Spondylitis Disease Activity Index; BASFI, Bath Ankylosing Spondylitis Functional Index; CAL, calprotectin; CDAI, Clinical Disease Activity Index; CRP, C-reactive protein; DAPSA, Disease Activity index for Psoriatic Arthritis; DAS28-CRP, Disease Activity Score in 28 joints with CRP; IRD, inflammatory rheumatic diseases; PhGA, physician global assessment; PGA, patient global assessment; RA, rheumatoid arthritis; axSpA, axial spondyloarthritis; PsA, psoriatic arthritis.

### Elevated serum calprotectin identifies patients with higher disease activity and more severe rheumatoid arthritis

We next evaluated the ability of sCAL to identify patients with active disease. The discriminatory performance of sCAL for polyarticular disease, defined as the presence of SJC≥4, was superior to that of CRP (AUC 0.67 versus 0.62, Delong p=0.012; Figure 3A). Using the predefined threshold of 4.2 μg/mL, sCAL identified patients with extensive joint involvement (SJC28≥4) with a specificity of 90%, although sensitivity remained poor (30.5%). In comparison, CRP ≥5 mg/L demonstrated lower specificity (76.8%) and a sensitivity of 42.7%. The performance of sCAL and CRP for identifying patients with moderate to high disease activity (CDAI >10) did not differ significantly (Figure 3B). Patients with elevated sCAL exhibited consistently higher disease activity across all clinical and composite disease activity measures (Figure 3C).

**Figure 3.**
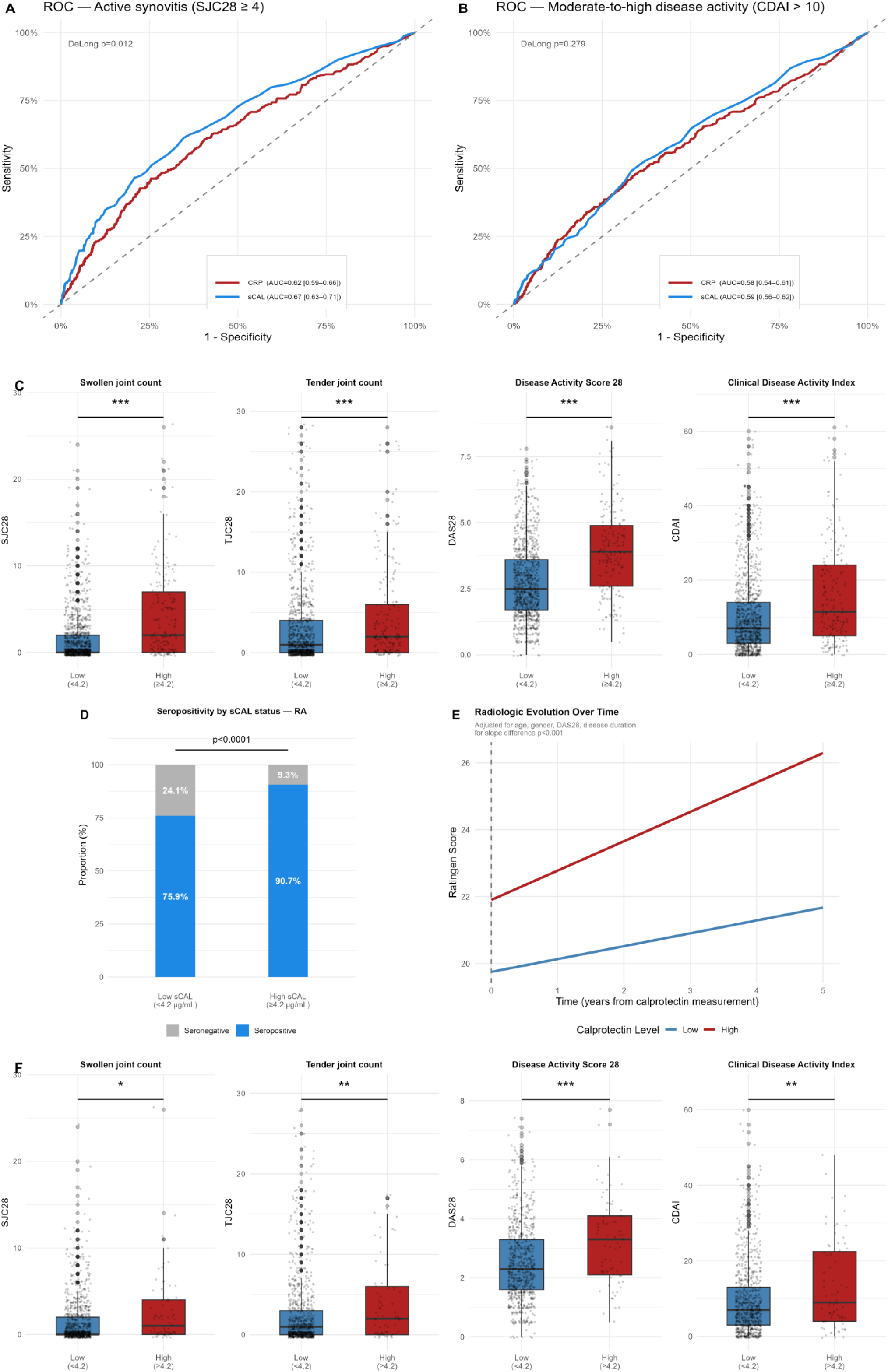
Elevated sCAL predicts increased disease activity and severity in rheumatoid arthritis. **(A)** Receiver Operating Characteristic (ROC) curves for sCAL and CRP to predict active synovitis (SJC28≥4) in RA. sCAL demonstrated superior discriminative performance compared to CRP (AUC 0.67 vs 0.62, DeLong p=0.012). **(B)** ROC curves for sCAL and CRP to predict moderate-to-high disease activity (CDAI>10). AUC comparison by DeLong method. **(C)** Disease activity scores stratified by sCAL status (Low <4.2 µg/mL vs High ≥4.2 µg/mL). Boxes indicate median and IQR. Wilcoxon rank-sum test. ***p<0.001. **(D)** Proportion of seropositive patients by sCAL status. Numbers indicate proportions within each group. Chi-square test, p<0.0001. **(E)** Radiological progression of joint damage (Ratingen score) over time stratified by sCAL status. Lines represent predicted values from a linear mixed-effects model adjusted for age, gender, DAS28, and disease duration. Patients with High sCAL exhibit significantly faster radiological progression compared to Low sCAL patients (p for slope difference=9.56×10⁻⁴). **(F)** Disease activity scores in RA patients with normal CRP (<5 mg/L, n=1067), stratified by sCAL status. Higher sCAL was associated with greater CDAI, DAS28, SJC28, and TJC28 (all p<0.05).

We further investigated the relationship between sCAL and established markers of RA severity. Elevated sCAL was strongly associated with RA-associated autoimmunity, with 90.7% of patients in the high-sCAL group being seropositive, compared with 75.9% in the low-sCAL group (p < 0.001; Figure 3D). Regarding articular damage progression, elevated sCAL was associated with accelerated radiographic progression over time, as assessed by the Ratingen score (Figure 3E). This association remained significant after adjustment for age, sex, DAS28 and disease duration (p<0.001). Collectively, these findings demonstrate that elevated sCAL identifies a subset of RA patients characterized by higher inflammatory activity, greater autoantibody positivity, and increased risk of structural disease progression. Moreover, the persistence of these associations in patients with normal CRP suggests that sCAL provides complementary prognostic information beyond conventional inflammatory biomarkers.

### sCAL discriminates disease activity independently of IL-6 pathway blockade

Importantly, associations between sCAL and disease severity remained significant among patients with normal CRP levels (CRP <5 mg/L; n = 1067) (Figure 3F). These findings suggest that sCAL captures clinically relevant inflammatory burden beyond that reflected by CRP alone. Given that CRP synthesis is directly regulated by IL-6, we examined whether sCAL retained its ability to reflect disease activity in patients treated with IL-6 inhibitors, in whom CRP is pharmacologically suppressed. Among RA patients with elevated sCAL (≥4,2µg/mL), those receiving anti-IL-6R therapy exhibited higher clinical disease activity than those receiving anti-TNF (CDAI, p<0.05), with a similar trend for SJC (p=0.071), whereas CRP did not differ between these groups (Supplementary figure 2A). These findings indicate that sCAL captures inflammatory activity independent of IL-6 signaling, providing complementary information in clinical settings where CRP may be unreliable.

### Elevated serum calprotectin is independently associated with increased mortality and cardiovascular risk

Given the established links between calprotectin and CV diseases, we next investigated the association between sCAL and adverse CV outcomes in patients with RA. Patients with elevated sCAL levels exhibited significantly reduced overall survival compared with those with low sCAL concentrations (log-rank p < 0.0001; Figure 4A). In multivariable Cox regression adjusting for potential confounders for cardiovascular risk, elevated sCAL remained independently associated with increased mortality risk. The magnitude of this association varied according to the cause of death, with an adjusted hazard ratios of 3.0 [1.87-4.78] for all-cause mortality (p<0.001), 5.9 [1.97-17.62] for respiratory-related mortality (p=0.002), and up to 7.5 for CV mortality (p=0.01). In contrast, no significant association was observed between elevated sCAL and infection-related mortality. Elevated sCAL was also associated with an increased risk of incident MACE. Survival analyses demonstrated a significantly higher cumulative incidence of MACE among patients with elevated sCAL levels (log-rank p=0.01; Figure 4B), and multivariable analyses identified high sCAL as an independent predictor of incident MACE, with an adjusted hazard ratio of 2.1 [1.044-4.244] compared with patients with low sCAL concentrations (Figure 4C). Importantly, these associations were also observed in the RA patient group alone. Elevated sCAL remained an independent predictor of all-cause mortality after adjustment for relevant clinical covariates (HR 1.49, [1.14–1.93]; p = 0.003; Supplementary Figure 3). Other independent predictors of mortality in RA included older age, male sex, diabetes mellitus, and glucocorticoid use. This association remained unchanged after additional adjustment for smoking status (HR 1.49 [1.15–1.94]; p = 0.003), confirming that the prognostic value of sCAL is independent of tobacco exposure.

**Figure 4.**
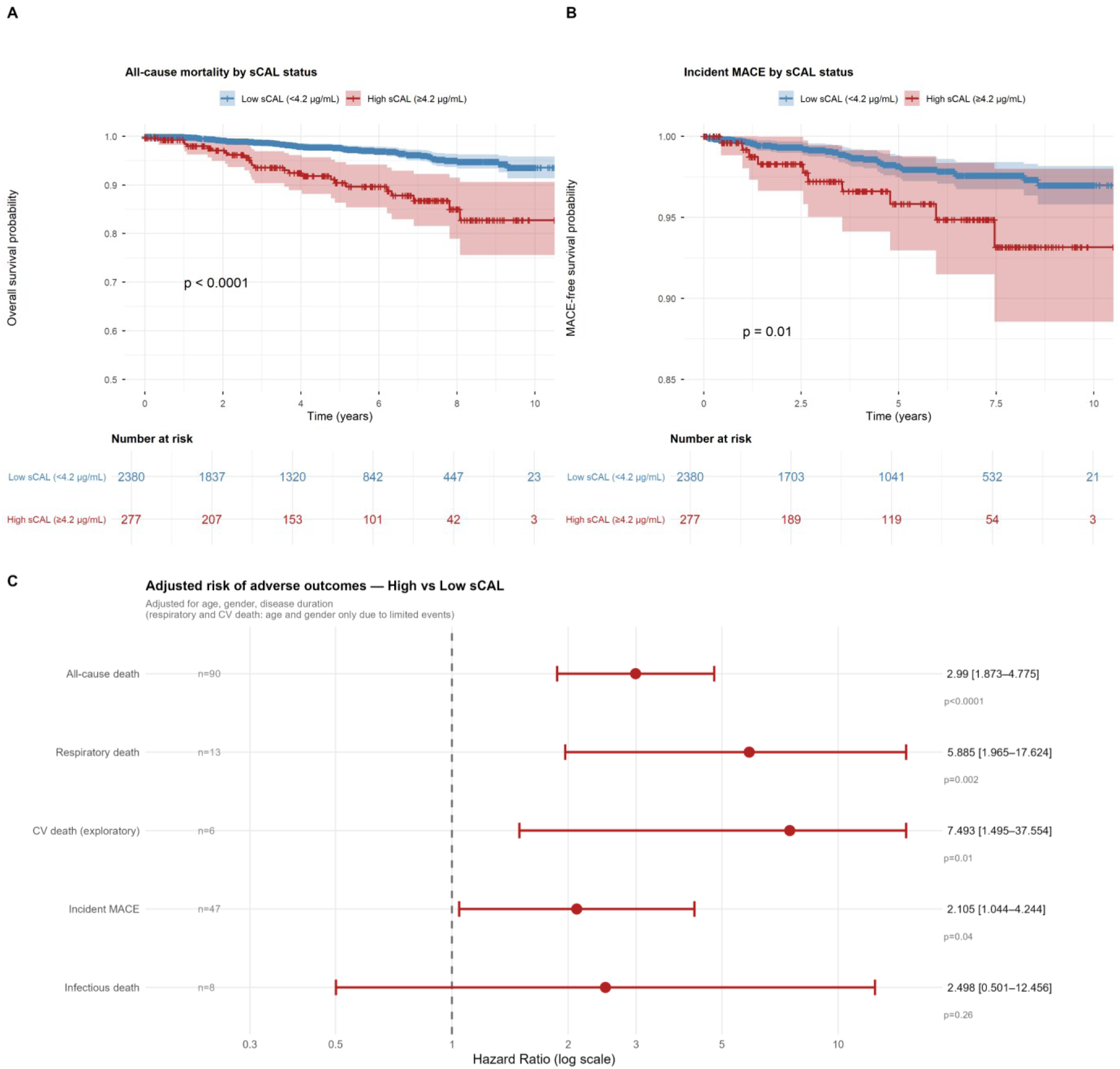
Elevated sCAL predicts increased mortality and cardiovascular events. **(A)** Kaplan-Meier survival curves for all-cause mortality stratified by sCAL status (Low <4.2 µg/mL vs High ≥4.2 µg/mL). Log-rank test p<0.0001. **(B)** Kaplan-Meier curves for incident major adverse cardiovascular events (MACE) by sCAL status. Log-rank test p=0.01. Numbers at risk are shown below each plot. **(C)** Forest plot of adjusted hazard ratios (HR) for adverse outcomes comparing High vs Low sCAL. Cox proportional hazards models adjusted for age, gender, and disease duration. Respiratory and CV death models adjusted for age and gender only due to limited events (n=13 and n=6 respectively). Arrows indicate confidence intervals extending beyond the axis limit. CV death is shown as exploratory only.

To further explore the relationship between sCAL and CV risk, we examined associations with established cardiovascular biomarkers. Across all three disease groups, sCAL was positively correlated with both hs-cTnT and NT-proBNP, two established markers of myocardial injury and cardiac dysfunction (Figure 5A-C). Elevated sCAL was independently associated with increased NT-proBNP concentrations after adjustment for age, sex, and underlying disease, with an adjusted odds ratio of 1.2 (p=0.008) (Figure 5B). Notably, these associations were independent of lipid profile, as sCAL did not correlate significantly with HDL-cholesterol, LDL-cholesterol or triglycerides (not shown). Patients with high sCAL (≥4.2 µg/mL) showed significantly elevated hs-cTnT and NT-proBNP levels compared to those with low sCAL (Figure 5C).

**Figure 5.**
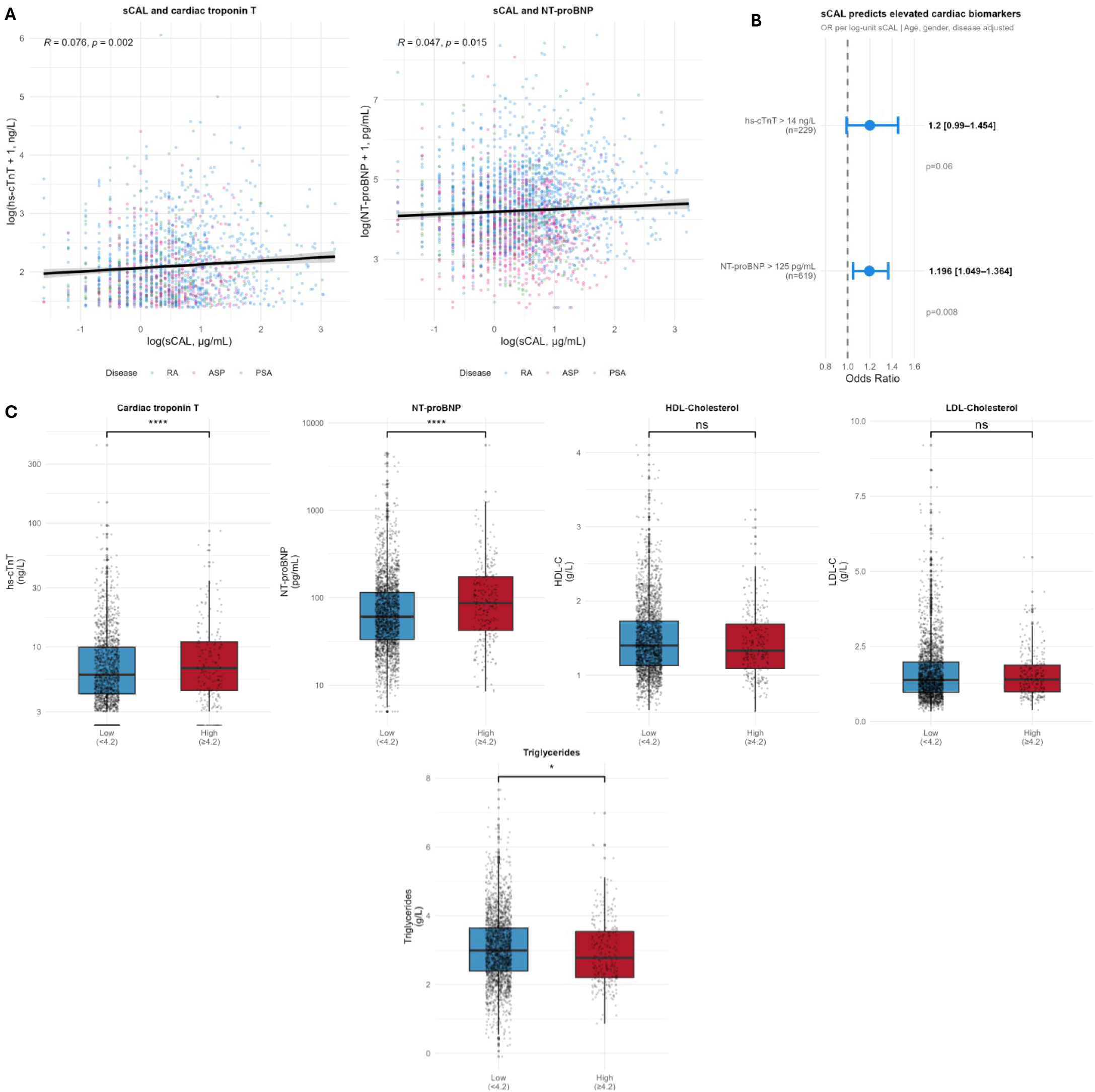
Elevated serum calprotectin identifies subclinical myocardial injury independently of lipid profile in inflammatory rheumatic diseases. **(A)** Association between sCAL and cardiac biomarkers. Left and middle panels show Spearman correlations between log-transformed sCAL and cardiac biomarkers by disease group. Right panel shows odds ratios for elevated cardiac biomarkers per log-unit increase in sCAL, adjusted for age, gender, and disease. hs-cTnT threshold: 14 ng/L (ESC 2019 Universal Definition of Myocardial Infarction). NT-proBNP threshold: 125 pg/mL (ESC Heart Failure Guidelines 2021). **(B)** Cardiovascular biomarkers stratified by sCAL status (Low <4.2 µg/mL vs High ≥4.2 µg/mL). Wilcoxon rank-sum test. ***p<0.001, *p<0.05, ns: not significant.

Collectively, these findings identify elevated sCAL as an independent marker of adverse prognosis in inflammatory rheumatic diseases, associated not only with increased mortality and CV events, but also with biomarkers of subclinical CV injury and dysfunction.

## Discussion

In this large real-world cohort of patients with IRDs, we demonstrate that elevated sCAL is preferentially associated with active RA, more so than with AS and PsA. Elevated sCAL levels were associated with increased disease activity, greater disease severity, accelerated structural damage progression, and adverse cardiovascular outcomes. Collectively, these findings support sCAL as a clinically relevant biomarker that captures both inflammatory burden and long-term prognosis in RA.

Although sCAL concentrations were elevated across inflammatory diseases, associations with disease activity were strongest in RA, where sCAL correlated consistently with SJC and composite disease activity indices. Conversely, associations in axSpA and PsA were weaker and generally inferior to those observed with CRP. These findings suggest that circulating calprotectin predominantly reflects the synovial inflammatory processes characteristic of RA and may therefore represent a disease-specific marker of peripheral joint inflammation. However, the diagnostic performance of sCAL for distinguishing RA from other IRD remained modest. By comparison, sCAL has shown substantial diagnostic utility in other inflammatory rheumatic diseases, particularly adult-onset Still’s disease [16].

Beyond its association with disease activity, sCAL also emerged as a marker of disease severity. Using a threshold of 4.2 μg/mL derived from a healthy blood donor population, we identified a subgroup of RA patients characterized by greater inflammatory activity, higher seropositivity rates, and more rapid structural joint damage progression. Taken together, these findings indicate that the principal value of sCAL in RA lies in disease stratification and prognostication rather than diagnostic discrimination. A notable finding of this study is the performance of sCAL compared with CRP. Across multiple measures of disease activity, sCAL performed at least as well as, if not better than CRP. Furthermore, elevated sCAL remained associated with increased disease activity even among patients with normal CRP concentrations. This observation is clinically relevant, as a substantial proportion of patients with active IRD do not exhibit elevated CRP levels despite ongoing inflammatory activity [6]. In such patients, sCAL may provide complementary information and could represent a valuable alternative marker for monitoring disease activity [17]. This complementarity was particularly evident in patients treated with IL-6 inhibitors, in whom CRP is pharmacologically suppressed through direct inhibition of hepatic synthesis. In this setting, sCAL, a marker of neutrophil-derived and local inflammation, remained associated with clinical disease activity despite IL-6 receptor blockade, whereas CRP did not. As calprotectin is released directly by activated neutrophils and monocytes [18] and is less dependent on hepatic IL-6-driven acute-phase responses than CRP, it may provide complementary information for monitoring residual inflammation when conventional acute-phase reactants are pharmacologically suppressed [17,19].

Our study also highlights a strong relationship between sCAL and CV risk. Cardiovascular disease is a major contributor to morbidity and mortality in RA, where chronic inflammation accelerates atherosclerosis and promotes cardiovascular dysfunction [20,21]. Elevated sCAL was independently associated with all-cause mortality, CV mortality, respiratory mortality, and major adverse CV events. The association between sCAL and all-cause of mortality was more modest in the RA subgroup (HR 1.49) than in the overall cohort (HR 3.0), likely reflecting the higher baseline cardiovascular burden in RA and the additional adjustment for the use of glucocorticoids and other treatments. In addition, sCAL correlated with established markers of CV injury and dysfunction, including high-sensitivity cardiac troponin T and NT-proBNP. These findings support the concept that calprotectin reflects inflammatory pathways involved not only in joint disease but also in systemic vascular and cardiac injury [22,23]. These associations suggest that sCAL may contribute to improved CV risk stratification in patients with RA and other IRDs, consistent with growing evidence implicating calprotectin in CV inflammation and atherosclerosis [24]. Calprotectin is expressed by activated myeloid cells within vascular lesions and has been associated with coronary artery disease, adverse CV outcomes, and CV mortality, supporting its potential role as a biomarker linking systemic inflammation to CV risk [25]. To our knowledge, no previous study has comprehensively evaluated the association between serum calprotectin and long-term CV outcomes in a large real-world RA cohort.

Several limitations should be acknowledged. Firstly, the analyses were performed within a real-world registry, resulting in heterogeneity in disease duration, treatment exposure, and follow-up schedules. Although elevated sCAL was independently associated with all-cause mortality, this finding should be interpreted in light of a limitation of the SCQM cohort, namely the incomplete follow-up and the lack of systematic ascertainment of occurrence of death, which may have led to underreporting. Nevertheless, significant associations were consistently observed despite this heterogeneity, supporting the robustness and generalizability of the findings. Secondly, sCAL was measured at a single time point, preventing assessment of longitudinal biomarker trajectories and their relationship with treatment response. Finally, although multiple confounders were accounted for in adjusted analyses, residual confounding cannot be excluded.

Overall, the present findings provide a strong rationale for further investigation of sCAL as a prognostic biomarker in RA. Prospective studies are required to determine whether serial sCAL measurements improve prediction of disease progression, treatment response, and long-term outcomes beyond conventional inflammatory markers. Particular attention should be given to complications that contribute substantially to RA-related mortality, including cardiovascular disease and RA-associated interstitial lung disease. Future studies should also evaluate whether incorporation of sCAL into routine disease assessment improves risk stratification and clinical decision-making.

## Conclusion

In conclusion, sCAL is a clinically relevant biomarker in RA that reflects both disease activity and disease severity. Its discriminative performance is comparable to, and in several analyses, superior to CRP, including among patients with normal CRP concentrations, where conventional inflammatory markers may fail to capture residual disease activity. This advantage was particularly apparent in patients treated with IL-6 inhibitors, in whom CRP is pharmacologically suppressed, highlighting a clinical setting where sCAL may be especially informative. Elevated sCAL independently identifies patients at increased risk of structural progression, cardiovascular events, and mortality.

These findings support the integration of sCAL into future strategies for disease monitoring, risk stratification, and prognostic assessment in RA.

## Acknowledgements

The authors are grateful to BÜHLMANN Laboratories AG for providing sCAL reagents free of charge. BÜHLMANN Laboratories AG had no involvement in the analysis or interpretation of the results. The authors also thank Per Riegler, Catherine Raptis for their valuable contributions and assistance.

The authors thank Dr.Pierre Lescuyer and his team at the serum repository (sérothèque) of the Geneva University Hospital (HUG) for providing access to the biological samples used in this study.

A list of rheumatology offices and hospitals contributing to the SCQM registry can be found on https://www.scqm.ch/en/aboutscqm/active-institutions/. The SCQM thanks the patients for their participation in the registry.

## Data availability statement

This study is based on data collected by the SCQM Foundation (Swiss Clinical Quality Management in Rheumatic Diseases), which operates a nationwide registry for rheumatic diseases. Access to the data is subject to restrictions and requires approval from the SCQM Foundation in accordance with the SCQM Rules of Research and Collaboration (https://www.scqm.ch/en/research/research-with-scqm-data/). Interested parties may contact the SCQM Foundation to request access to the data for research purposes.

