## supplementary Tables and Figures for "Serum Calprotectin as predictor of Rheumatoid Arthritis disease severity and Mortality"

**Suppl table 1. Baseline characteristics of axSpA patients stratified by sCAL status**

| Variable | Overall (n=771) | Low sCAL (<4.2 µg/mL, n=725) | High sCAL (≥4.2 µg/mL, n=46) | p-value |
|---|---|---|---|---|
| **Demographics** | | | | |
| Age, years | 43.10 [34.70, 52.00] | 43.50 [34.92, 52.10] | 37.00 [29.73, 48.20] | 0.005 |
| Male sex, n (%) | 452 (58.6) | 418 (57.9) | 32 (69.6) | 0.160 |
| Smoker, n (%) | 462 (59.9) | 427 (59.1) | 32 (69.6) | 0.214 |
| Disease duration, years | 6.46 [2.51, 12.84] | 6.55 [2.67, 13.21] | 4.04 [0.52, 10.19] | 0.010 |
| **Inflammatory biomarkers** | | | | |
| CRP, mg/L | 1.84 [0.67, 5.50] | 1.70 [0.66, 4.71] | 7.05 [2.16, 30.08] | <0.001 |
| sCAL, µg/mL | 1.50 [1.00, 2.40] | 1.40 [0.90, 2.20] | 5.05 [4.70, 5.77] | <0.001 |
| **Disease activity** | | | | |
| BASDAI | 3.80 [1.80, 5.65] | 3.80 [1.80, 5.60] | 4.80 [2.20, 6.00] | 0.253 |
| ASDAS | 2.30 [1.50, 3.20] | 2.20 [1.50, 3.10] | 2.90 [2.00, 3.80] | 0.004 |
| BASFI | 1.90 [0.50, 4.45] | 1.80 [0.50, 4.40] | 2.60 [0.88, 5.12] | 0.316 |
| **Disease manifestations*** | | | | |
| Sacroiliitis, n (%) | 553 (80.4) | 516 (80.0) | 34 (85.0) | 0.571 |
| Enthesitis, n (%) | 450 (66.2) | 426 (66.8) | 21 (53.8) | 0.139 |
| Dactylitis, n (%) | 65 (9.6) | 59 (9.3) | 5 (13.5) | 0.569 |
| Coxitis, n (%) | 134 (19.9) | 123 (19.4) | 11 (28.2) | 0.261 |
| Skin manifestation, n (%) | 49 (6.9) | 44 (6.6) | 5 (11.9) | 0.323 |
| **Comorbidities** | | | | |
| Hypertension, n (%) | 125 (16.2) | 118 (16.3) | 6 (13.0) | 0.702 |
| Diabetes, n (%) | 46 (6.0) | 45 (6.2) | 1 (2.2) | 0.421 |
| **Cardiovascular biomarkers** | | | | |
| hs-cTnT, ng/L | 3.19 [0.00, 5.45] | 3.16 [0.00, 5.37] | 3.28 [0.00, 5.72] | 0.684 |
| NT-proBNP, pg/mL | 43.65 [25.10, 80.35] | 43.80 [25.70, 79.90] | 39.80 [23.80, 86.25] | 0.746 |
| **Treatments** | | | | |
| Glucocorticoids, n (%) | 12 (1.6) | 11 (1.5) | 1 (2.2) | 1.000 |
| csDMARDs, n (%) | 226 (29.3) | 212 (29.4) | 14 (30.4) | 1.000 |
| Statins, n (%) | 62 (8.0) | 60 (8.3) | 2 (4.3) | 0.498 |
| Current bDMARD, n (%) | | | | <0.001 |
| None | 254 (32.9) | 233 (32.3) | 20 (43.5) | |
| Anti-TNF | 496 (64.3) | 474 (65.7) | 20 (43.5) | |
| Abatacept | 1 (0.1) | 1 (0.1) | 0 (0.0) | |
| Anti-IL6 | 5 (0.6) | 3 (0.4) | 2 (4.3) | |
| Other | 15 (1.9) | 11 (1.5) | 4 (8.7) | |
| **Outcomes** | | | | |
| Prevalent MACE, n (%) | 27 (3.5) | 26 (3.6) | 1 (2.2) | 0.923 |
| Incident MACE, n (%) | 5 (0.6) | 4 (0.6) | 1 (2.2) | 0.705 |
| All-cause death, n (%) | 5 (0.6) | 4 (0.6) | 1 (2.2) | 0.705 |

*Data are presented as median [interquartile range] for continuous variables and n (%) for categorical variables. p-values: Wilcoxon rank-sum test (continuous); Chi-squared or Fisher's exact test (categorical).*
**Disease manifestations coded as yes/no/unknown; unknown treated as missing.*
*Caution: High sCAL group n=46 — results should be interpreted with caution due to limited sample size.*
*Abbreviation: ASDAS, Ankylosing Spondylitis Disease Activity Score; axSpA, Axial spondyloarthritis; BASDAI, Bath Ankylosing Spondylitis Disease Activity Index; BASFI, Bath Ankylosing Spondylitis Functional Index; bDMARD, biologic disease-modifying antirheumatic drug; CRP, C-reactive protein; csDMARD, conventional synthetic DMARD; hs-cTnT, high-sensitivity cardiac troponin T; MACE, major adverse cardiovascular event; NT-proBNP, N-terminal pro-B-type natriuretic peptide; sCAL, serum calprotectin.*

**Suppl table 1. Baseline characteristics of PsA patients stratified by sCAL status**

| Variable | Overall (n=434) | Low sCAL (<4.2 µg/mL, n=412) | High sCAL (≥4.2 µg/mL, n=22) | p-value |
|---|---|---|---|---|
| **Demographics** | | | | |
| Age, years | 51.85 [43.15, 59.50] | 51.85 [43.15, 59.65] | 50.00 [41.65, 55.10] | 0.291 |
| Male sex, n (%) | 245 (56.5) | 235 (57.3) | 9 (40.9) | 0.197 |
| Smoker, n (%) | 120 (27.6) | 109 (26.6) | 11 (50.0) | 0.032 |
| Disease duration, years | 5.29 [2.34, 10.80] | 5.37 [2.36, 10.93] | 4.26 [0.86, 9.91] | 0.324 |
| **Inflammatory biomarkers** | | | | |
| CRP, mg/L | 1.72 [0.75, 4.38] | 1.69 [0.73, 4.15] | 10.07 [2.18, 26.77] | <0.001 |
| sCAL, µg/mL | 1.40 [0.90, 2.10] | 1.40 [0.80, 1.90] | 4.80 [4.45, 6.20] | <0.001 |
| **Disease activity** | | | | |
| DAS28 | 2.20 [1.40, 3.10] | 2.20 [1.40, 2.95] | 3.80 [2.80, 4.95] | <0.001 |
| CDAI | 7.00 [3.00, 12.00] | 6.00 [3.00, 12.00] | 14.00 [3.75, 23.00] | 0.032 |
| Swollen joint count (28) | 0.00 [0.00, 1.00] | 0.00 [0.00, 1.00] | 1.00 [0.00, 4.50] | 0.008 |
| Tender joint count (28) | 0.00 [0.00, 2.00] | 0.00 [0.00, 2.00] | 2.00 [0.50, 6.00] | 0.006 |
| **Disease manifestations (PSA-specific)*** | | | | |
| Skin manifestation, n (%) | 345 (89.1) | 327 (89.3) | 17 (85.0) | 0.811 |
| Dactylitis, n (%) | 178 (48.1) | 165 (47.1) | 12 (63.2) | 0.261 |
| Enthesitis, n (%) | 218 (59.9) | 208 (60.6) | 10 (50.0) | 0.478 |
| Nail manifestations, n (%) | 182 (50.3) | 172 (50.1) | 9 (50.0) | 1.000 |
| Sacroiliitis, n (%) | 85 (28.7) | 83 (29.2) | 2 (18.2) | 0.650 |
| Coxitis, n (%) | 39 (11.0) | 38 (11.4) | 1 (5.6) | 0.703 |
| **Comorbidities** | | | | |
| Hypertension, n (%) | 102 (23.5) | 97 (23.7) | 4 (18.2) | 0.739 |
| Diabetes, n (%) | 33 (7.6) | 30 (7.3) | 3 (13.6) | 0.500 |
| **Cardiovascular biomarkers** | | | | |
| hs-cTnT, ng/L | 3.92 [0.00, 6.52] | 3.88 [0.00, 6.51] | 4.85 [0.00, 6.28] | 0.509 |
| NT-proBNP, pg/mL | 52.10 [30.23, 87.42] | 51.15 [29.98, 84.73] | 91.00 [33.25, 140.75] | 0.066 |
| **Treatments** | | | | |
| Glucocorticoids, n (%) | 22 (5.1) | 19 (4.6) | 3 (13.6) | 0.170 |
| csDMARDs, n (%) | 272 (62.7) | 260 (63.4) | 11 (50.0) | 0.298 |
| Statins, n (%) | 44 (10.1) | 42 (10.2) | 2 (9.1) | 1.000 |
| Current bDMARD, n (%) | | | | 0.949 |
| None | 148 (34.1) | 141 (34.4) | 6 (27.3) | |
| Anti-TNF | 227 (52.3) | 213 (52.0) | 13 (59.1) | |
| Abatacept | 3 (0.7) | 3 (0.7) | 0 (0.0) | |
| Anti-IL6 | 4 (0.9) | 4 (1.0) | 0 (0.0) | |
| JAK inhibitor | 3 (0.7) | 3 (0.7) | 0 (0.0) | |
| Other | 49 (11.3) | 46 (11.2) | 3 (13.6) | |
| **Outcomes** | | | | |
| Prevalent MACE, n (%) | 19 (4.4) | 19 (4.6) | 0 (0.0) | 0.618 |
| Incident MACE, n (%) | 7 (1.6) | 7 (1.7) | 0 (0.0) | 1.000 |
| All-cause death, n (%) | 7 (1.6) | 6 (1.5) | 1 (4.5) | 0.804 |

*Data are presented as median [interquartile range] for continuous variables and n (%) for categorical variables. p-values: Wilcoxon rank-sum test (continuous); Chi-squared or Fisher's exact test (categorical).*
**Disease manifestations coded as yes/no/unknown; unknown treated as missing.*
*Caution: High sCAL group n=22 — results should be interpreted with caution due to very limited sample size.*
*Abbreviation: bDMARD, biologic disease-modifying antirheumatic drug; CDAI, Clinical Disease Activity Index; CRP, C-reactive protein; csDMARD, conventional synthetic DMARD; DAS28, Disease Activity Score 28 joints; hs-cTnT, high-sensitivity cardiac troponin T; MACE, major adverse cardiovascular event; NT-proBNP, N-terminal pro-B-type natriuretic peptide; PsA, psoriatic arthritis; sCAL, serum calprotectin.*

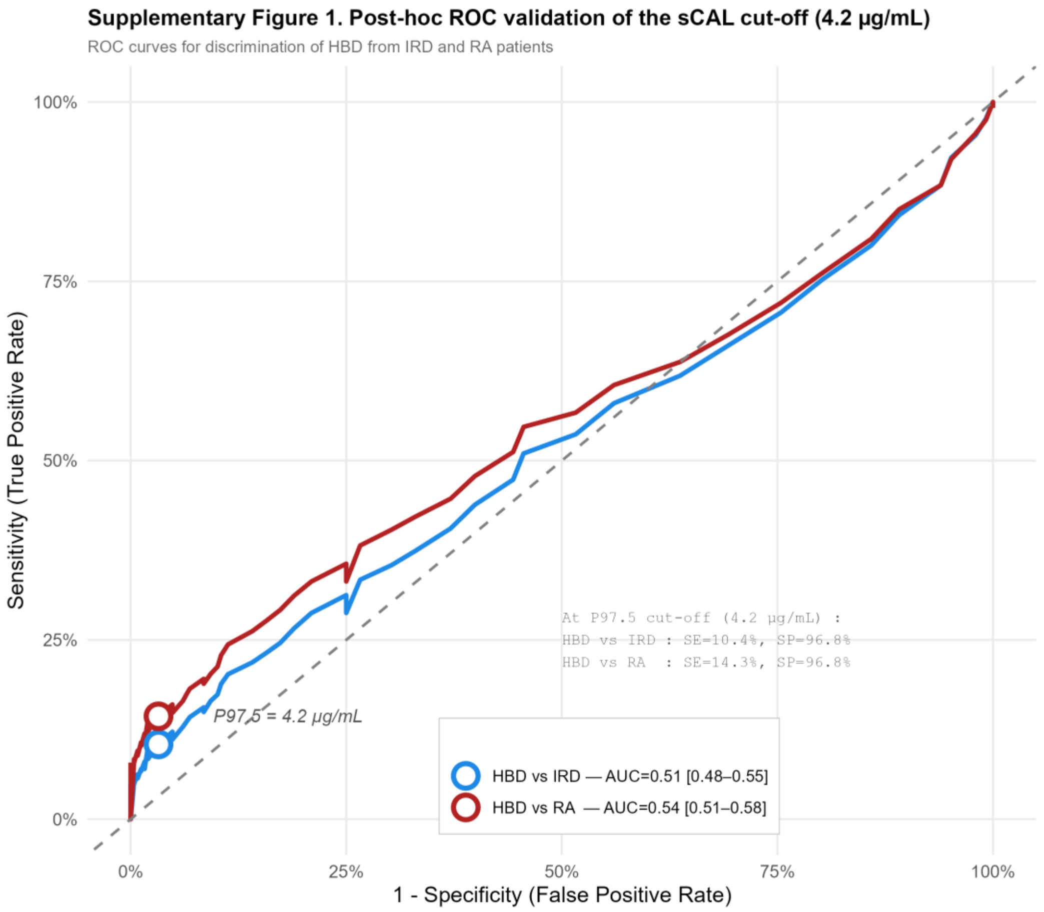


**Supplementary figure 1**. **Post-hoc ROC validation of the sCAL cut-off (4.2ug/mL).** Receiver operating characteristic (ROC) curves for the discrimination of healthy blood donors (HBD) from patients with inflammatory rheumatic diseases (IRD, blue) and from patients with rheumatoid arthritis (RA, red). Open circles indicate the operating point at 4.2ug/mL.

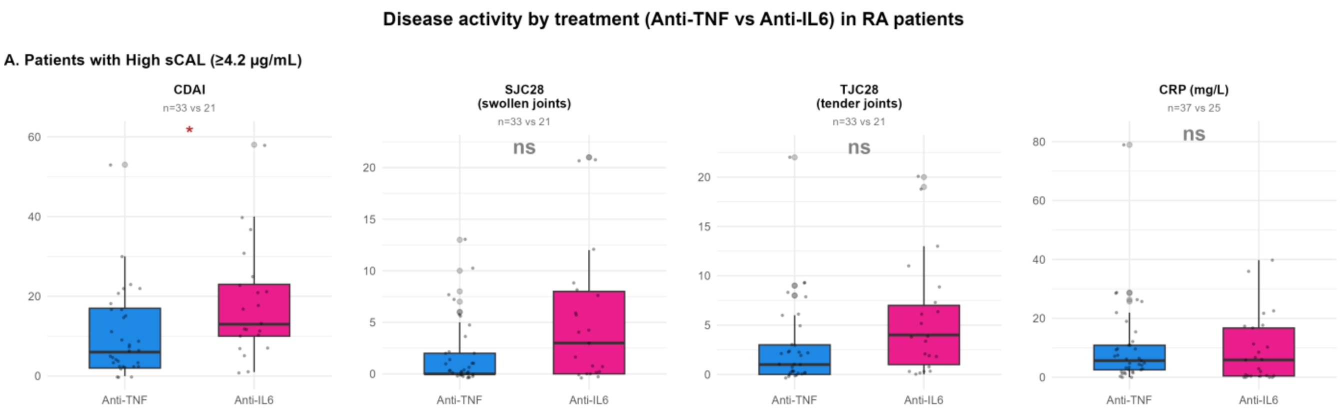


**Supplementary figure 2**. **Serum calprotectin reflects disease activity independently of IL-6 pathway blockade in rheumatoid arthritis.** Comparison of disease activity measures and inflammatory biomarkers between RA patients treated with anti-TNF versus anti-IL-6 therapy, stratified by biomarker status. Among patients with elevated sCAL (≥4.2 µg/mL), clinical disease activity (CDAI, swollen joint count [SJC28], tender joint count [TJC28]) and CRP were compared between treatment groups. Sample sizes are indicated below each subpanel. Statistical comparisons were performed using the Wilcoxon rank-sum test. *p<0.05, **p<0.01, ***p<0.001; ns, not significant.

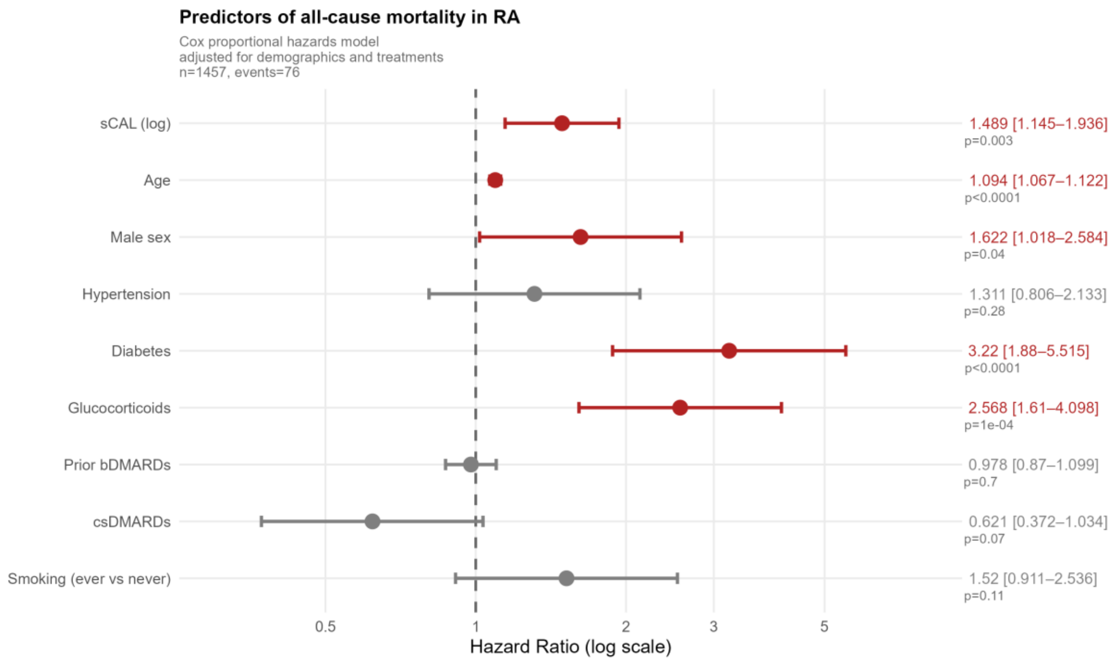


**Supplementary figure 3**. **Independent predictors of all-cause mortality in rheumatoid arthritis.** Forest plot of adjusted hazard ratios (HR) with 95% confidence intervals from a multivariable Cox proportional hazards model for all-cause mortality in RA patients (n=1,457; 76 events). The model was adjusted for demographics and treatments, including log-transformed sCAL, age, sex, hypertension, diabetes, glucocorticoid use, prior bDMARD exposure, and csDMARD use. Red indicates statistically significant predictors (p<0.05); grey indicates non-significant variables. The dashed vertical line denotes HR=1 (no effect). Hazard ratios are displayed on a logarithmic scale.
